# Unveiling Local Radiation Events through Metagenome Assembled Genomes of Archaea and Bacteria in Hypersaline Microbial Mats from the Archean Domes Site, Cuatro Cienegas, Coahuila, Mexico

**DOI:** 10.1101/2023.08.19.553462

**Authors:** Ulises Erick E Rodríguez Cruz, Hugo G Castelán-Sánchez, David Madrigal-Trejo, Luis Enrique Eguiarte, Valeria Souza

## Abstract

A comprehensive study was conducted in the Cuatro Cienegas Basin (CCB) in Coahuila, Mexico, known for its remarkable microbiological diversity and unique physicochemical properties. The ″Archaean Domes ″ (DA) in the CCB are hypersaline, non-lithifying microbial mats. This study focused on analyzing the small domes and circular structures formed in DA through metagenome assembly genomes (MAGs) with the aim of finding new microorganisms and providing information on the tree of life in a place as diverse as the CCB. In total, 329 MAGs were identified, including 52 archaea and 277 bacteria. Remarkably, 30 Archaea and 154 Bacteria could not be classified at the genus level, highlighting the remarkable diversity of CCB. The CCBs showed significant diversity at the phylum level, with Proteobacteria being the most abundant, followed by Euryarchaeota, Firmicutes, Bacteroidetes, Actinobacteria, Cyanobacteria, Spirochaetes, Chloroflexi, Planctomycetes, Candidatus Parvarchaeota, Verrucomicrobia, Balneolaeota, Nitrospirae, and Tenericutes. Subsequently, the MAGs were classified into a phylogenetic tree. In Archaea, monophyletic groups MAGs belonged to the phyla Archaeoglobi, Candidatus Aenigmarchaeota, Candidatus Nanoarchaeota, Candidatus Lokiarchaeota, and Halobacteriota. Among the Bacteria, monophyletic groups were identified as well, including Spirochaetes, Proteobacteria, Planctomycetota, Actinobacteria, Verrucomicrobiota, Bacteroidetes, Bipolaricaulota, Desulfobacterota, and Cyanobacteria. These monophyletic clusters may indicate radiation events that are likely influenced by geographical isolation as well as extreme environmental conditions reported in AD pond like phosphorus deficiency (122:42:1 C:N:P), fluctuating pH and a salinity of 5.28%

## 1.0 Introduction

In recent years, culture-independent techniques have revolutionized our understanding of microbial diversity and evolutionary relationships within the tree of life. Notably, the discovery of the Candidate Phylum Radiation (CPR group) by Hug et al. (2016) and the novel archaeal phylum Lokiarchaeota by Spang et al. (2015) have had profound implications for our knowledge of microbial taxonomy, which have significantly expanded the taxonomic coverage of the tree of life. For example, Gong et al. (2022) identified several novel bacterial phyla within the FCB superphylum, highlighting the importance of MAGs in uncovering previously unknown microbial lineages and their ecological roles.

The Cuatro Cienegas Basin (CCB) in the Chihuahua Desert of northern Mexico has been the subject of extensive studies that have discovered novel lineages due to its unique physicochemical properties; the nitrogen-phosphorus (N:P) ratio in the CCB can vary widely, from conditions of severe phosphorus deficiency to near-normal conditions, affecting microbial proliferation (Elser et al., 2000). This environment has triggered evolutionary responses in endemic microorganisms, such as the reduction of the genome of Bacillus coahuilensis and its production of sulfolipids instead of phospholipids as potential adaptations to low phosphorus concentrations (Bonilla-Rosso et al., 2012; Souza et al., 2008; Alcaraz et al., 2008).

Previous metagenomic diversity profiles in two different microbial mats in CCB (red mat and green mat) showed that bacteria dominated in the red mat with a relative abundance of 98% with the most abundant phyla being Proteobacteria, Firmicutes, and Cyanobacteria, while Archaea and Eukarya represented only 1.78% 0.26%, respectively. Similarly, at the green mat site, a relative abundance of ∼93% was found for bacteria (without a dominant phylum), only 2.06% for Archaea, and 2.79% for Eukaryota (Bonilla-Rosso et al. 2012). On the other hand, using 16S rRNA gene tags, 5,167 OTUs (with 97% identity) were detected in soil, sediment, and water samples at different sampling sites in the Churince system (now defunct hydrological system within the CCB). This diversity represented 60 different phyla of microorganisms, of which only three belonged to the Archaea (Souza et al., 2018). Overall, despite its high bacterial diversity, the Archaea domain is underrepresented at several sites in the CCB.

However, this changed in March 2016 when an unexpected rain exposed a shallow pond (see Figure 1) on the Pozas Azules ranch of Pronatura Noroeste within the CCB (26° 49’ 41.7″ N, 102° 01’ 28.7″ W). Dome-shaped structures formed around orange circles; these structures are only observed under humid conditions after heavy rain, at most once a year. Inside these elastic dome structures, an anoxic, carbon dioxide and methane-rich interior has been observed which reminisce what the atmosphere could have been during most of the Archean eon, before the oxygenation of the atmospheres and oceans, hence the name ″Archean domes ″ (DA) has been used to describe this site (Espinosa-Asuar et al., 2022; Medina-Ch&aacute;vez et al., 2023; Madrigal-Trejo et al., 2023). A pH of 9.8 and a salinity of 5.28% were measured during the rainy season, while in the dry season the pH is 5 and the salinity reaches saturation. During the rainy season, a stoichiometric imbalance of C:N:P of 122:42:1 has been reported (Espinosa-Asuar et al., 2022; Medina-Ch&aacute;vez et al., 2019; Madrigal-Trejo et al., 2023).

**Figure 1.**
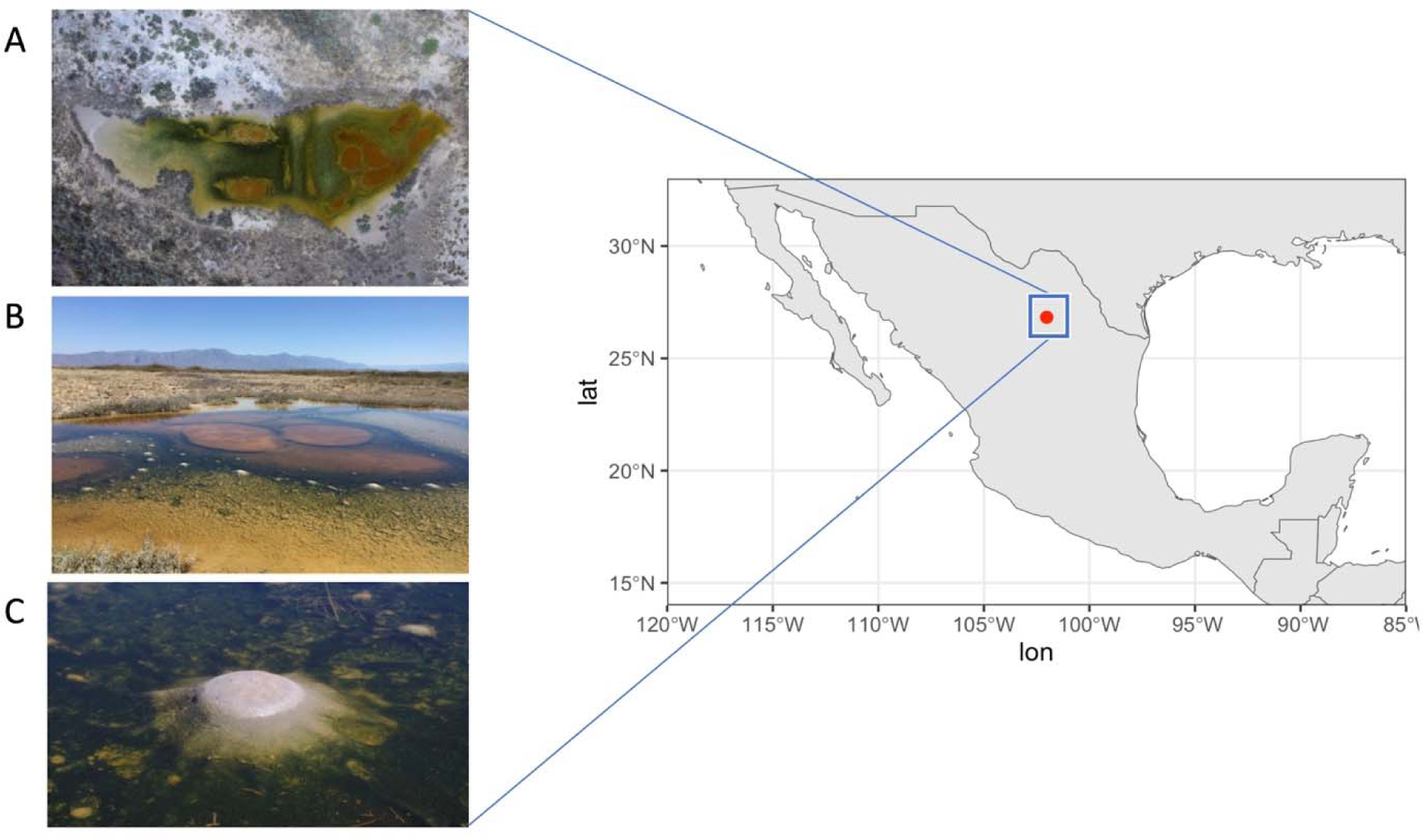
Archean Domes (AD) site. Panel A displays an aerial photograph of the site, while Panel B exhibits an image taken in 2016, the year of the site’s initial study. The orange circles indicate the characteristic areas of the site that were measured, along with dome-shaped structures, one of which is depicted in greater detail in Panel C. Photo credit: David Jaramillo.

In an initial study of microbial mat diversity in the AD site by Medina-Ch&aacute;vez et al. (2023) revealed that the relative abundance of the Archaea domain was up to ∼5%, corresponding to 5 Archaea phyla, 25 orders, 36 families, 93 genera and 230 species (with threshold of 97%). Compared to the diversity analysis conducted at other sites in CCB where the relative abundance of Archaea barely reached 2.0% (Bonilla-Rosso et al., 2012; Souza et al., 2018). Subsequent studies from the AD site reported that in 2019 there was a significant increase in the relative abundance of Archaea of ∼30.60%, with the phylum Euryarchaeota being the most abundant (Madrigal-Trejo et al., 2023).

Considering that the CCB has a high level of endemism and AD is particularly diverse, we expected to find many novel lineages with the help of MAGs. Therefore, in this study, we aimed to analyze MAGs in AD ponds in two areas: the zone of dome formation and orange circles. We reconstructed the genomes and placed them in a phylogenetic context for taxonomic classification. Our analysis yielded a comprehensive set of 329 MAGs from the AD system, encompassing both archaeal and bacterial domains. Among these, we identified 52 MAGs belonging to the Archaea domain and 277 MAGs belonging to the Bacteria domain. These genomes covered a remarkable range of new microorganisms, spanning 56 phyla in both domains.

## 2.0 Material and methods

### 2.1 Sample Collection, Genomic DNA Extraction and Sequencing

Samples were collected using a soil corer with 8 PVC tubes to obtain replicates for each sample. Fragments of ∼10 cm deep were collected from the surface area of domes between 2016 and 2019, and starting in October 2020, sampling was expanded to include domes and circles at different depths, down to ∼50 cm below the surface. Each sample was promptly transferred into liquid nitrogen for preservation and stored until DNA extraction was performed using the MP FastDNA™ SPIN kit for Soil. Information regarding the sample depth and year of collection is provided in **Table 1**, it is important to note that in 2019 and 2020, sampling was conducted during two different seasons (wet and dry seasons) of the same year.

**Table 1.**
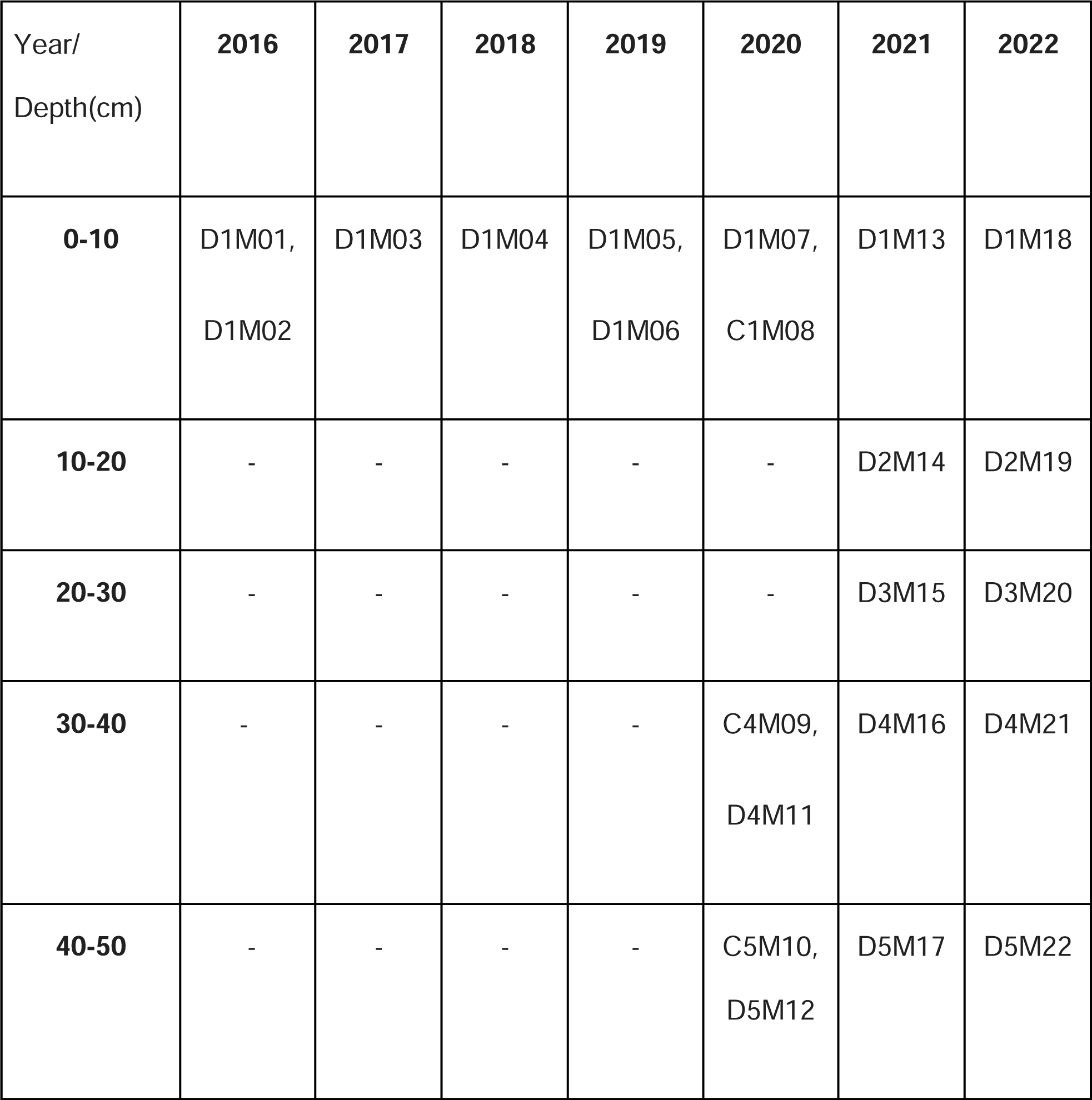
Codes of the samples and seasonal variations in the sampling of the CCB Domes (D) and Circles (C).

| Year/<br>Depth(cm) | 2016 | 2017 | 2018 | 2019 | 2020 | 2021 | 2022 |
| --- | --- | --- | --- | --- | --- | --- | --- |
| **0-10** | D1M01,<br>D1M02 | D1M03 | D1M04 | D1M05,<br>D1M06 | D1M07,<br>C1M08 | D1M13 | D1M18 |
| **10-20** | - | - | - | - | - | D2M14 | D2M19 |
| **20-30** | - | - | - | - | - | D3M15 | D3M20 |
| **30-40** | - | - | - | - | C4M09,<br>D4M11 | D4M16 | D4M21 |
| **40-50** | - | - | - | - | C5M10,<br>D5M12 | D5M17 | D5M22 |

Samples were sequenced in Cinvestav-Langebio (http://langebio.cinvestav.mx/labsergen/) Irapuato, Mexico, using Miseq Reagent Kit v3 2×300 pb paired-end on the Ilumina Miseq platform.

### 2.2 Metagenomic Analysis

We quality filtered the raw sequencing data using FastQC (v0.11.8) (Andrews, S. 2010) and Trimmomatic (v0.39) (Bolger, Lohse & Usadel, 2014). Reads were then taxonomically classified using Kaiju (v1.8.1) (Menzel, Ng & Krogh, 2016) using default parameters. Results were visualized using R (v4.1.0) (R Core Team, 2021) with ggplot2 (v3.3.5) (Wickham H, 2016).

To determine whether there was a correlation between the relative abundance at the phylum level and the sampling depth, the Pearson correlation test was performed, which is implemented in R (v4.1.0). Assembly of reads was performed using MetaSPAdes (v3.15.3) (Nurk, Meleshko, Korobeynikov & Pevzner, 2017). The assembled contigs were required for binning, which was performed using MaxBin2 (v2.2.7) (Wu, Simmons & Singer, 2015) and MetaBat2 (v2.12.1) (Kang et al., 2019). To reduce contamination in the bins, the software Binning refiner (v1.4.2) (Song & Thomas, 2017) was used. MAGs integrity was assessed using CheckM (v1.1.3) (Parks et al., 2015) with default settings. Good quality MAGs were considered to have >70% completeness and <10% contamination.

### 2.3 Phylogenetic placement of MAGs

For taxonomic assignment and placement of MAGs in the phylogenetic tree of life, we used the GTDB-tk (v1.6.0) software toolkit (Chaumeil, Mussig, Hugenholtz & Parks, 2019), which identifies 122 and 120 archaeal and bacterial marker genes respectively, using HMMER (Eddy, 2011). Briefly, genomes were assigned to the domain with the highest proportion of identified marker genes. The selected domain-specific markers were aligned using HMMER, concatenated into a single multiple sequence alignment, and trimmed with the bacterial or archaeal ∼5000 column mask used by GTDB (Chaumeil, Mussig, Hugenholtz & Parks, 2019). This and the following phylogenetic trees were visualized with the iTOL tool (Letunic & Bork, 2021).

In addition to the previous phylogenetic taxonomic classification of MAGs, we performed a species tree encompassing all MAGs, both Archaea and Bacteria, to provide a general overview of the MAGs of the AD site. To achieve the above, the Orthofinder program (Emms & Kelly, 2019) was used, which implements the DIAMOND program for the inference of orthologous genes, MCL for the clustering algorithm, ETE Tree library for all tree management, MAFFT for multiple sequence alignment and FastTree for phylogenetic building. the phylogenomic tree resulting from this analysis is shown in **Figure S1**.

### 2.4 Validation of the monophyletic clades of radiation events of CCB MAGs

The taxonomic classification provided by GTDB-tk and observing patterns in the clustering of MAGs within the reference trees of this program suggest radiations (Radiation means the formation of a monophyletic taxon of at least three species by two or more speciation events (Sudhaus, 2004).

To validate the occurrence of radiation events, we investigated four distinct taxonomic groups (see Table S1 to S5 for additional genome information). The first group examined was the superphylum Asgardarchaeota, as it has been proposed to have a potential relationship with the origin of eukaryotes (Devos, 2021). The second group consisted of the Candidate phylum Bipolaricaulota, characterized by its deep branching within the bacterial domain and the presence of members capable of autotrophic carbon fixation via the ancient WL-pathway for carbon fixation (Takami et al., 2012). We further explored The Planctomycetes, Verrucomicrobia, Chlamydiae (PVC) superphylum, which encompasses bacteria exhibiting eukaryote-like cellular compartmentalization and varying degrees of cell organization (Kamneva et al., 2012). Lastly, we focused on Cyanobacteria due to their pivotal role as primary producers in the microbial ecosystem, influencing carbon and nitrogen cycles (Elster & Kv&iacute;derov&aacute;, 2011). Phylogenomic analyses were performed individually for each taxonomic group to investigate potential radiation events and evaluate the clustering patterns of Metagenome Assembled Genomes (MAGs). The main objective of these analyses was to determine if the MAGs within each taxonomic group demonstrate a monophyletic clustering pattern, which could indicate radiation events. This study aimed to shed light on the clustering behavior and evolutionary relationships of the MAGs within their respective taxonomic groups. To achieve this, we examined the phylogenetic relationships among the MAGs using both the MAGs obtained from the four taxonomic groups and genomes available in existing databases.

## 3.0 Results

### 3.1 Taxonomic profile of sequencing reads

In the taxonomic assignment of the sequencing reads (Figure 2, panels A and B), the 15 most abundant phyla in the 22 metagenomes are Proteobacteria (34.4%), follow by Euryarchaeota (14.45%), Firmicutes (11.31%), Bacteroidetes (11.02%), Actinobacteria (9.06%), Cyanobacteria (8.9%), Spirochaetes (2.03%), Chloroflexi (1.14%), Planctomycetes (0.99%), Candidatus Parvarchaeota (0.92%), Verrucomicrobia (0.78%), Balneolaeota (0.61%), Nitrospirae (0.42%) and Tenericutes (0.38%).

**Figure 2.**
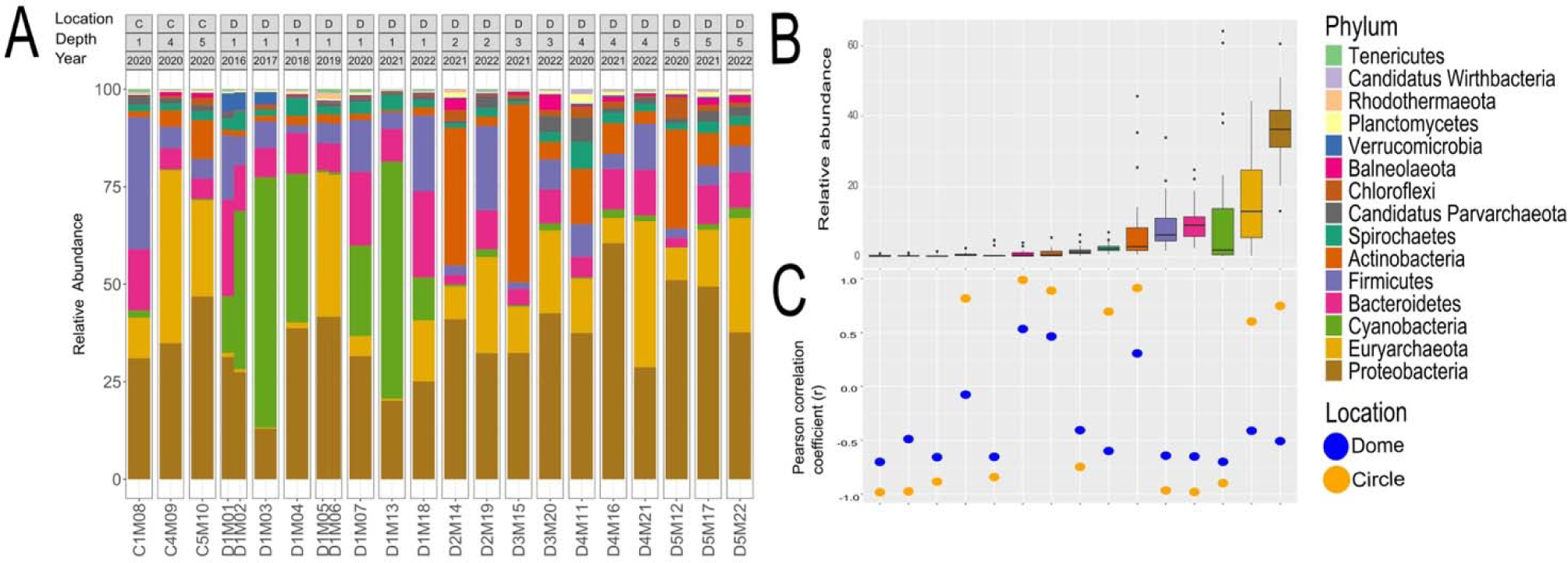
Taxonomic assignment of DA site sequencing reads. (A) Taxonomic classification with Kaiju at the phyla level using metagenomic reads. Labels above correspond to sampling area circle C) or dome D) and sampling depth (0-10 cm (1) to 40-50 cm (5)), and sampling year 2016 to 2022. B) The relative abundance of the most representative phyla. C) The values of Pearson correlation coefficient (r) between relative abundance and sampling depth.

Although Euryarchaeota is the only archaeal phylum with a high relative abundance, seven phyla (Thaumarchaeota, Candidatus Nanohaloarchaeota, Candidatus Lokiarchaeota, Crenarchaeota, Candidatus Korarchaeota, Candidatus Micrarchaeota, Nanoarchaeota) were found within the archaea domain with relative abundance values below 1.0%.

The correlation analysis between the sampling depth and relative abundance at phylum level (Figure 2C) showed that some phyla are found preferentially at the surface, such as Cyanobacteria, Bacteroidetes, Firmicutes, Parvarchaeota, Verrucomicrobiota, Rhodothermaeota, Wirdthbacteria and Tenericutes. Likewise, other phyla such as Balneolaeota, Chloroflexi and Actinobacteria, are found preferentially in deeper samples.

### 3.2 Metagenome-assembled genome quality

In this study, we are reporting a total of 329 MAGs, for Archaea (52) and Bacteria (277) (refer to Figure 3a and 3b and Table S5 for further information), all of them with an integrity percentage greater than 70% and contamination of less than 10%. For the archaeal MAGs, the genome size ranged from 556,584 pb to 4,091,783 pb (Table S5). Estimates for contamination of MAGs ranged from 0% to 5.61%. The N50 value ranged from 4,688 to 130,076 pb. The GC content has a wide range from 22.80% to 70.80%.

**Figure 3.**
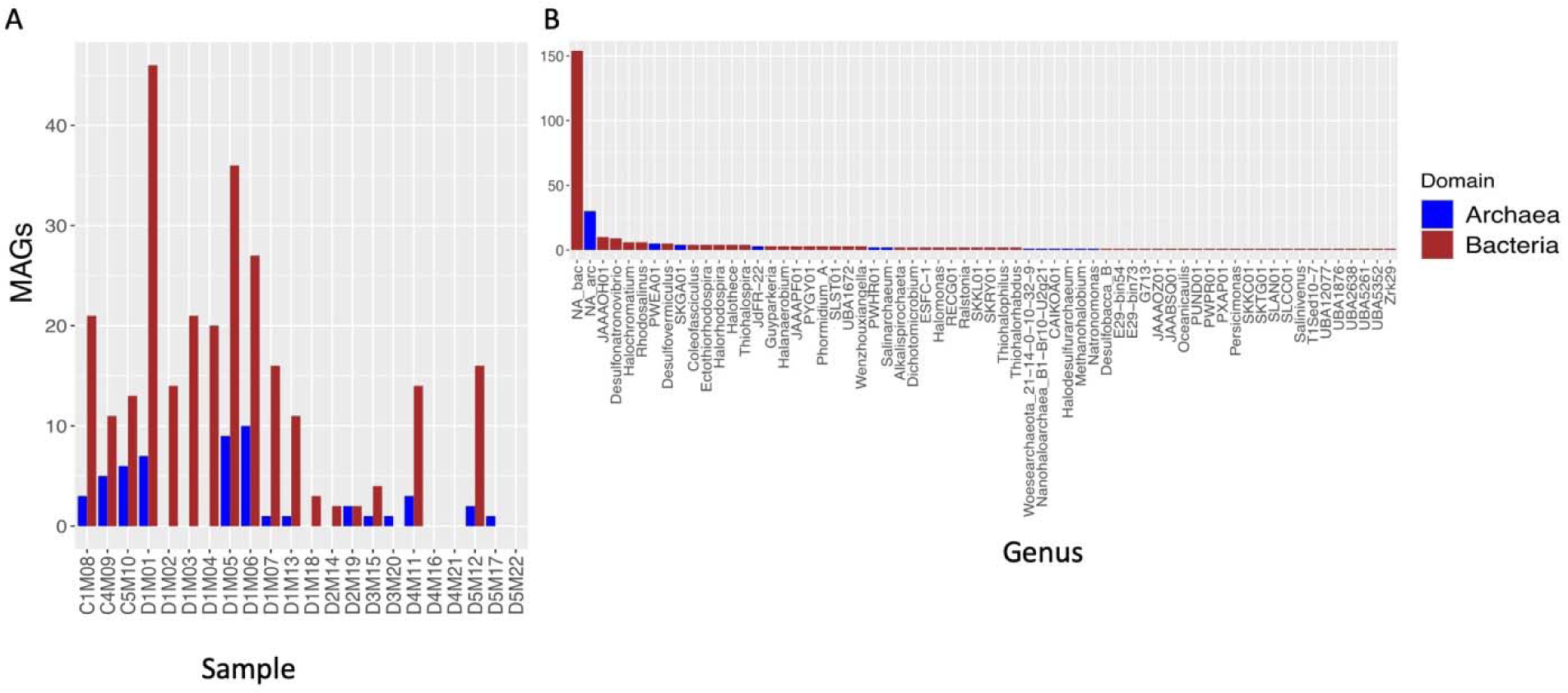
Summary of MAGs. (A) Number of MAGs obtained per sample. (B) MAG classified at the genus level, while most MAGs did not have a classification assigned to the genus level.

The quality scores for the bacterial MAGs ranged from 645,757 to 7,241,784 bp of genome size. The N50 value ranged from 5.317 to 128.031, and the GC content ranged from 29.7% to 73.73%. Most of the MAGs assembled over the years in all metagenomes correspond to bacteria, which could indicate a predominance of this domain in the mats of AD in CCB, while MAGs corresponding to archaea were found in a lower proportion.

The abundance of MAGs obtained varied depending on the depth of sampling. The highest number of MAGs (246) was obtained at a depth of about 0-10 cm, while only 6 MAGs were obtained at depths of about 10-20 cm and 6 MAGs for 20-30 cm. However, at depths of about 30-40 cm, 32 MAGs were recovered, and at depths of about 40-50 cm, 39 MAGs were recovered.

### 3.3 Taxonomic assignment and phylogenetic placement of MAGs

The 52 MAGs assigned to the Archaea domain were taxonomically divided into twelve classes, while the 277 MAGs of the Bacteria domain belong to 47 different taxonomic classes. Figure 4a shows the Archaea tree derived from GTDB-tk software to show the distribution of Archaea MAGs from a phylogenetic perspective.

**Figure 4.**
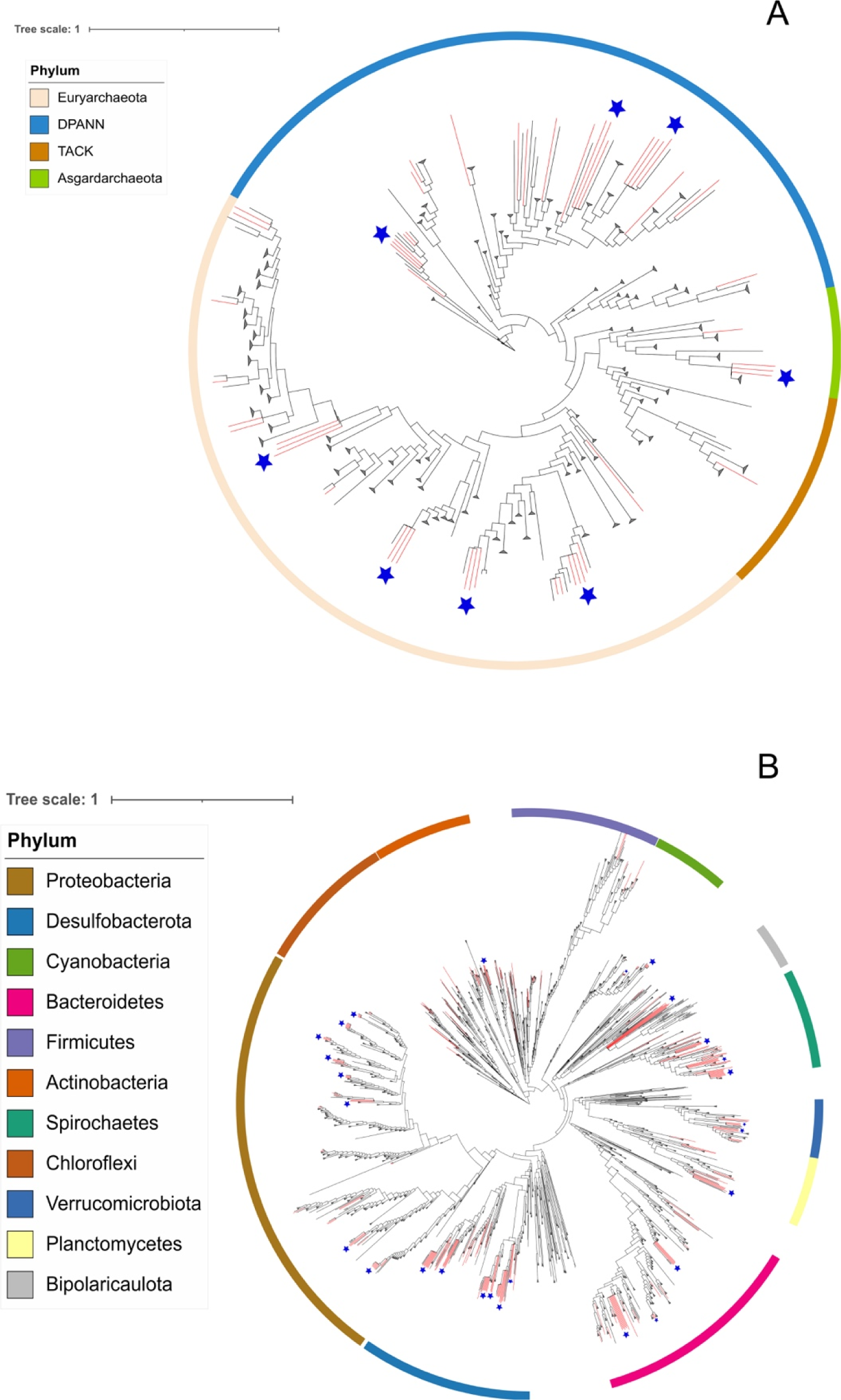
Phylogenomic placement of MAGs from the Archean Domes on the Archaea (A) and Bacteria (B) domain tree. The phylogenomic tree was reconstructed using GTDB-tk. The branches in red show the retrieved AD lineages.

The MAGs from AD were distributed across all three Archaea superphyla: DPANN, Asgard, TACK and phylum Euryarchaeota. The MAGs of Archaea are distributed across 12 different taxonomic classes: Archaeoglobi (3), Bathyarchaeia (2), Hadarchaeia (1), Halobacteria (9), Lokiarchaeia (3), Methanosarcinia (1), Micrarchaeia (1), Nanoarchaeia (14), Nanosalinia (2), candidate class PWEA01 (6), Thermoplasmata (9), and Thorarchaeia (1).

In the case of Bacteria (Figure 4b), the most abundant MAGs were: Proteobacteria (55), Desulfobacterota (53), Spirochaetes (32), Bacteroidota (28), Cyanobacteria (13), Planctomycetota (12), Verrucomicrobiota (11). Other less common phyla are not shown in Figure 4b, such as Fibrobacterota, Patescibacteria, Hydrogenedentota, Eremiobacterota, Goldbacteria, Marinisomatota, Myxococcota, Omnitrophota, Synergistota, Thermotogota, Acidobacteria, Armatimonadota, Campylobacterota, Gemmatimonadota and Sumerlaeota.

### 3.4 Radiation events

Some MAGs both in Archaea and Bacteria domains, exhibited clustering patterns that may suggest local radiation events as can be seen in the phylogenetic trees obtained in the taxonomic assignment of the MAGs with GTDB-tk and which are indicated with red arrows in both phylogenetic trees. For instance, the MAGs of the following Archaeal phyla: Euriarchaeota, Candidatus Aenigmarchaeota, Candidatus Nanoarchaeota, Candidatus Lokiarchaeota, showed such a pattern (Figure 4a). Similarly, in the case of the Bacteria domain, the phyla that show groupings that suggest radiation are Actinobacteria, Cyanobacteria, Bipolaricaulota, Spirochaetes, Verrucomicrobiota, Planctomycetes, Bacteroidetes, Desulfobacterota, and Proteobacteria.

The previous results led us to carry out phylogenetic analyzes separately from the phyla that have already been mentioned in the methodology; From these analyses, it can be seen in the corresponding figure 5a that the MAGs corresponding to the phylum Lokiarchaeota are grouped in a monophyletic manner. In the case of the phylum Bipolaricaulota (Figure 5b), our results show the formation of a monophyletic group of 4 MAGs (out of a total of 7 MAGs). Likewise, in the analysis of the MAGs of the PVC superphyllum, it is observed that two monophyletic groups are formed (figure 5c), the first one of 7 MAGs, whose taxonomic classification corresponds to the Phycisphaerae class, the other monophyletic group of the PVC superphylum is made up of 11 MAGs of the phylum Verrucomicrobia, order Opituales. Finally, the analysis of the cyanobacterial MAGs resulted in 3 monophyletic groups (Figure 5d), the first of 3 MAGs from the genus Phormidium, the second of 4 MAGs from the genus Coleofasciculus and the last group of 4 MAGs from the genus Halothece.

**Figure 5.**
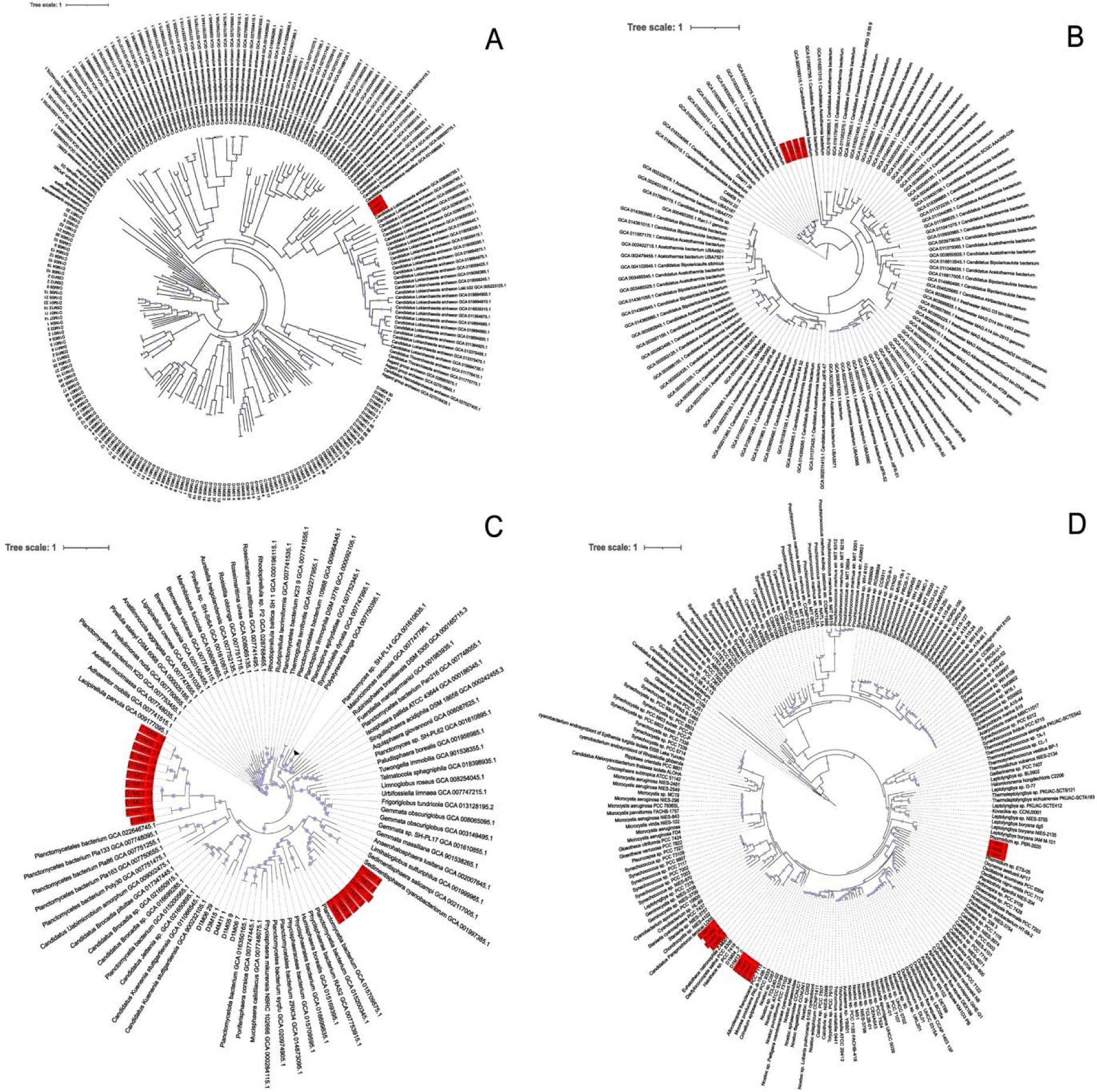
Phylogenetic trees of the Asgardarchaeota (A), Candidatus Bipolaricaulota (B), PVC (D) and Cyanobacteria (D) groups to validate radiation events. Highlighted in red are the leaves of individual trees representing MAGs from pnd AD, exhibiting monophyletic grouping.

The observations of the monophyletic groups in the phylogenetic trees of the 4 groups analyzed support our observation of the radiations in different phyla of the MAGs obtained at the DA site in CCB.

### 4.0 Discussion

This study aimed to recover MAG to provide information on the evolutionary relationships at the Archaean Domes site in the Cuatro Cienegas Basin (CCB). Unlike previous methods that used 16S rRNA tags or metagenome assemblies. For the first time in CCB, we used metagenome-assembled (MAG) genomes to expand our understanding of the phylogenetic tree of life. Our analysis revealed the presence of several predominant phyla, including Proteobacteria, Cyanobacteria, Firmicutes, Bacteroidetes, Actinobacteria, Spirochaetes, Chloroflexi, and Euryarchaeota, consistent with previous studies of diversity at the site (Espinosa Asuar et al., 2022; Madrigal-Trejo et al., 2023).

The archaeal community at the AD site exhibited monophyletic clades, indicating new archaea within AD, in contrast to other sites within the CCB where they are relatively rare, making AD an exceptional site for study of the Archeal domain of life.

Our analysis revealed that most of the genomes we present in this study correspond, at least at the species level, to previously unclassified genomes. This is not surprising considering that many of the phyla in which our MAGs are classified have only recently been described. For example, the Candidatus Aenigmarchaeota is an archaeal cluster first identified in 2013 as part of a study of ″microbial dark matter ″ (Rinke et al., 2013). The same is true for the Candidatus Woesearchaeota phylum and the Asgardarchaeota superphylum, which were also recently described (Castelle et al., 2015; Spang et al., 2015).

The Lokiarchaeota MAGs from AD show a monophyletic cluster pattern (see Figure 5a and S1), which may indicate that the Lokiarchaeota MAGs are endemic to the AD site. Whether the Lokiarchaeota of the CCB were cosmopolitan species, they would occur at multiple sites or have a non-monophyletic distribution scattered across different branches of the phylogenetic tree.

The diversity of the DA site in CCB is remarkable due to the distribution of the MAGs throughout almost the entire tree of life of Bacteria and Archaea. Compared to other hypersaline sites, only Shark Bay (blue hole mats) (Kindler et al., 2021) and Lake Hillier (Sierra et al., 2022) have reported members of the Asgard, TACK, DPANN and Euryarchaeota superphyla coexisting within the same microbial mat environment. This highlights the unique nature of the DA site and its potential as a source for studying the evolution and adaptation of microorganisms in extreme environments. The presence of Euryarchaeota, Asgard, and DPANN was reported in Guerrero Negro (Garc&iacute;a-Maldonado et al., 2022). The presence of Euryarchaeota, TACK, and DPANN has been reported in High-Bourne Cay (Khodadad & Foster, 2012). In Lake Magadi (Kambura et al., 2016) and Hamelin Pool (Ruvindy et al., 2015), only members of Euryarchaeota and TACK have been reported, whereas in Cape Recife (Waterworth et al., 2020), members of Asgard and DPANN have been reported. However, other analyses often report only the dominant relative abundance of some specific members of a single phylum, such as in Lake Tyrrel (Andrade et al., 2015), Euryarchaeota and Nanohaloarchaeota dominate in the Dead Sea (Jacob et al., 2017), and within the phylum Euryarchaeota, only members of of the family Halobacteriaceae. In Lake Meyghan (Naghoni et al., 2017), Euryarchaeota members dominate, and at the Hammam Essalihine site (Adjeroud et al., 2020), the Archaea domain is weakly represented by members of Parvarchaeota and Crenarchaeota, and at the Salar de Atacama sites (Laguna Brava and Tebenquiche) (Kurth et al., 2021), Lago Diamante (Rascovan et al., 2015), Socompa (Kurth et al., 2017), and Rottnest Island (Mendes Monteiro et al., 2020), only representatives of Euryarchaeota have been reported.

| Our study revealed the clustering of bacterial MAGs, indicating the occurrence of local radiation events. For instance, the AD site exhibited the presence of the newly proposed phylum Candidatus Bipolaricaulota (Hao et al., 2018), which showed monophyletic clustering. Similarly, our phylogenetic analyses of the Cyanobacteria and PVC group also demonstrated the formation of such monophyletic clusters. These findings suggest that the analyzed groups, as well as other groups displaying similar phylogenetic patterns, likely represent endemic groups. The observed clustering pattern aligns with the oligotrophic conditions characterized by limited phosphorus availability reported at the site, as evidenced by a reported C:N:P ratio of 122:42:1 (Espinosa-Asuar et al., 2022; Medina-Ch&aacute;vez et al., 2019; Madrigal-Trejo et al., 2023).

Previous studies have suggested that the low phosphorus (P) conditions in the CCB have triggered an evolutionary response among its endemic microorganisms, exemplified by *B. coahuilensis*. Remarkable adaptations to the environment have been observed in *B. coahuilensis*, including its ability to produce sulpholipids instead of phospholipids. This adaptation is attributed to the absence of genes responsible for synthesizing P-rich teichoic acids and polyanionic teichuronic acids (Souza et al., 2008).

Furthermore, our analysis identified MAGs from the phylum Spirochaetes, which were particularly abundant within the bacterial phyla. Notably, Spirochaetes were not previously detected in large numbers in other microbial diversity studies in CCB (Souza et al. 2018), and 32 MAGs were present in the AD site. We also identified MAGs from the versatile phylum Bacteroidetes which has been reported both in environmental samples and in the human gut, and furthermore, these bacteria are important and dominant carbohydrate degraders (Munoz et al., 2016).

In summary, our study provides information on the remarkable diversity and unique characteristics of microorganisms at site AD in the Cuatro Cienegas basin. Metagenome-assembled (MAG) genomes have allowed us to broaden our understanding of the tree of life and reveal the presence of a diverse microbial community, which, viewed from a phylogenetic perspective, suggests that the DA site might harbor many endemic lineages.

## 5.0 Conclusions

In this study, we present the discovery of 329 metagenome-assembled genomes (MAGs) from the DA site, comprising both Archaea (52) and Bacteria (277), encompassing a remarkable 56 phyla across both domains. The DA site is known for its fluctuating salinity and pH and has long been of interest because of its unique physicochemical conditions that support the growth of extremophile organisms.

Our findings underscore the exceptional microbiological diversity in this environment, as none of the MAGs could be classified at the species level and a significant portion (184 MAGs) could not be classified at the genus level. These results strongly suggest the presence of previously unknown microbial species and genera at this site. The collection of MAGs obtained in this study represents a valuable resource that can contribute to expanding our understanding of microbial diversity within the tree of life.

The DA site showcases remarkable microbiological diversity. However, phylogenetic analyses also unveil clustering patterns that suggest the occurrence of radiation events. Determining the nature of these events, whether adaptive or not, is beyond the scope of this article. Nonetheless, one of the main objectives of this article was to provide evidence supporting the existence of such radiation events.

## 6.0 Data availability statement

All data, including metagenome raw reads and metagenome assembled genomes, were deposited in the NCBI database in the bioproject PRJNA847603 with the accession identifiers SAMN35734740 to SAMN35735068

## 9.0 Acknowledgments

We would like to thank R. Tapia-Lopez, E. Aguirre-Planter and M. Rosas, for technical and field assistance. We also thank PRONATURA Noreste for the access to the Pozas Azules ranch, and to E. Rebollar-Caudillo and A.Becerra-Bracho for their valuable feedback on the manuscript. We specially want to thank R. Garcia-Herrera, for facilitating the use of high performace computing cluster ″Patung ″ located on Laboratorio Nacional de Ciencias de la Sostenibilidad (LANCIS), Instituto de Ecolog&iacute;a (UNAM).

---

**Figure S1.**
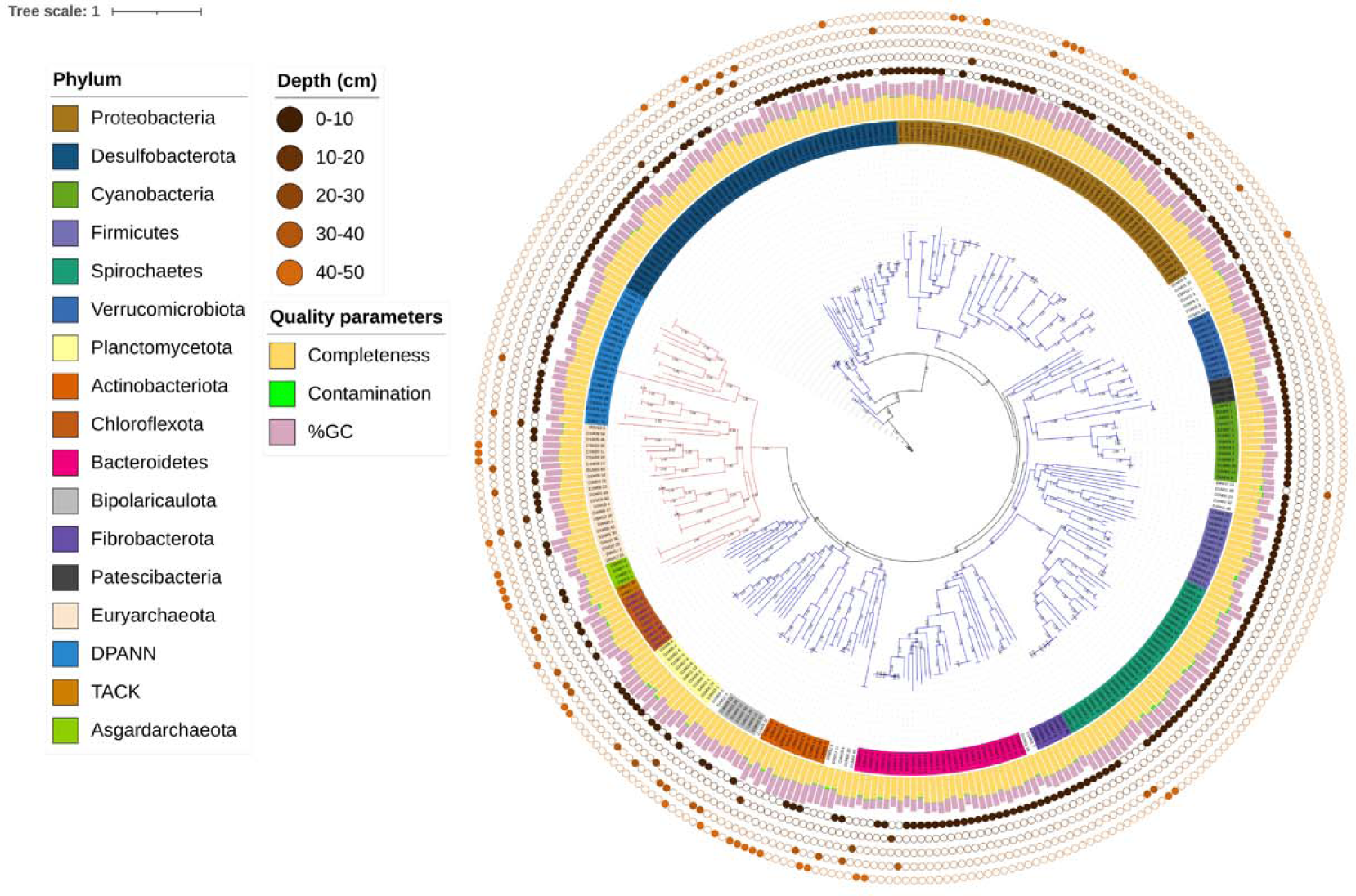
Phylogenomic tree including both Archaeal and Bacterial domain of the unique MAGs from AD site in CCB. The concentric circles with different dots show the sample origin, from surface to 40-50 cm. The bars show the values of contamination, completeness and %GC

